# MicroED data collection with SerialEM

**DOI:** 10.1101/446658

**Authors:** M. Jason de la Cruz, Michael W. Martynowycz, Johan Hattne, Tamir Gonen

## Abstract

We developed a procedure for the cryoEM method MicroED using SerialEM. With this approach, SerialEM coordinates stage rotation, microscope operation, and camera functions for automated continuous-rotation MicroED data collection. More than 300 datasets can be collected overnight in this way, facilitating high-throughput MicroED data collection for large-scale data analyses.

---

The cryoEM method MicroED is gaining momentum and is at the stage at which automation could further expand its use. In MicroED, a nanocrystal is exposed to the electron beam in diffraction mode and is continuously rotated while the diffraction pattern is collected on a fast camera as a movie^1,2^. This mode of MicroED data collection is called continuous rotation and is analogous to the rotation method in X-ray crystallography^3^. Combined with a camera operating in shutterless mode, continuous-rotation MicroED avoids partially-measured reflections by sampling all reciprocal space throughout the rotation range of the crystal, and provides both faster data collection and simpler data processing^1^. Continuous rotation has also improved the quality of the measurements^1^ and allowed MicroED to tackle more complex structure determination projects^4^; it has therefore become the standard method of data collection in MicroED. MicroED data are collected from a grid which might contain thousands of nanocrystals. A maximum of ~140° of data can be collected per crystal (+/- 70° tilt) so that, depending on symmetry, a single nanocrystal may yield an almost complete dataset^5^. Data from several crystals can be merged to increase completeness if each crystal is oriented differently on the grid^6^. MicroED data is then processed using standard X-ray crystallography software where structures are determined and refined as described before^7,8^.

Until now, all MicroED data were collected manually. Thus, the method could benefit from automation, where multiple targets are selected for acquisition in a single run, as is commonly done with other cryoEM modalities such as electron tomography (ET)^9^ and single particle analysis (SPA)^10^. These methodologies utilize software that controls the electron microscope and camera in order to perform the operations necessary during long-running data acquisition sessions with minimum human intervention. Automation is attractive because it increases the throughput of the instrument, which is particularly relevant to busy facilities where microscopes are operated as shared resources. It also reduces error arising from manual execution of repetitive tasks and allows for data collection 24/7.

SerialEM^11^ is a freely-available and open-source Microsoft Windows-based program used to coordinate microscope tasks and acquire digital images on TEMs. As of September 2018, SerialEM is installed on over 500 electron microscopes worldwide. It is compatible with modern electron microscopes and imaging detectors from several major manufacturers, and provides a consistent user interface across different hardware platforms. The software is highly extensible and allows microscope and camera acquisition tasks to be automated through its scripting command processor. Updates to the software and user-created scripts have enabled increasingly automated SPA^12^ and ET^13^ data collection. Other TEM automation software programs exist primarily for imaging mode only^10^ (*e.g*., for SPA and ET); SerialEM is currently the only one of these with the ability to control the microscope in diffraction mode in conjunction with a continuously-rotating stage.

We have developed a procedure using a script for SerialEM which enables large-scale MicroED data collection on TEMs by Thermo Fisher Scientific (formerly FEI Company, Philips Electron Optics) coupled to electron detectors from various manufacturers. An overview of this procedure is presented in **Figure 1** and generally follows a similar protocol commonly used for collecting SPA data. Once a grid containing nanocrystals is loaded into the microscope, the operator identifies potential crystals either by inspecting the grid manually using low-dose imaging^14^, or by montaging a whole-grid atlas. An atlas provides a fast way to visually screen an entire grid for crystals offline and keeps the exposure of the sample at a minimum. The location of each crystal is then marked in a similar way to selecting regions of holes/gridsquares for SPA data collection. The operator can then collect a sample diffraction pattern from each of the marked crystals to narrow down the selection of crystals for complete data collection (**Figure 2**). This step can be done either manually or automatically in SerialEM using its Navigator tools. Before data collection can begin on a selected set of crystals, the eucentric height needs to be determined at each data collection point on the grid. Akin to crystal centering in X-ray diffraction, this ensures that the crystal does not move out of the confines of the measurement during the continuous-rotation experiment^2^. Several reviews and papers have been published on crystal identification, troubleshooting, and microscope setup for diffraction^2,7,15–17^. The stage coordinates of the selected, well-diffracting crystals are then loaded to the automated data collection pipeline.

**Figure 1.**
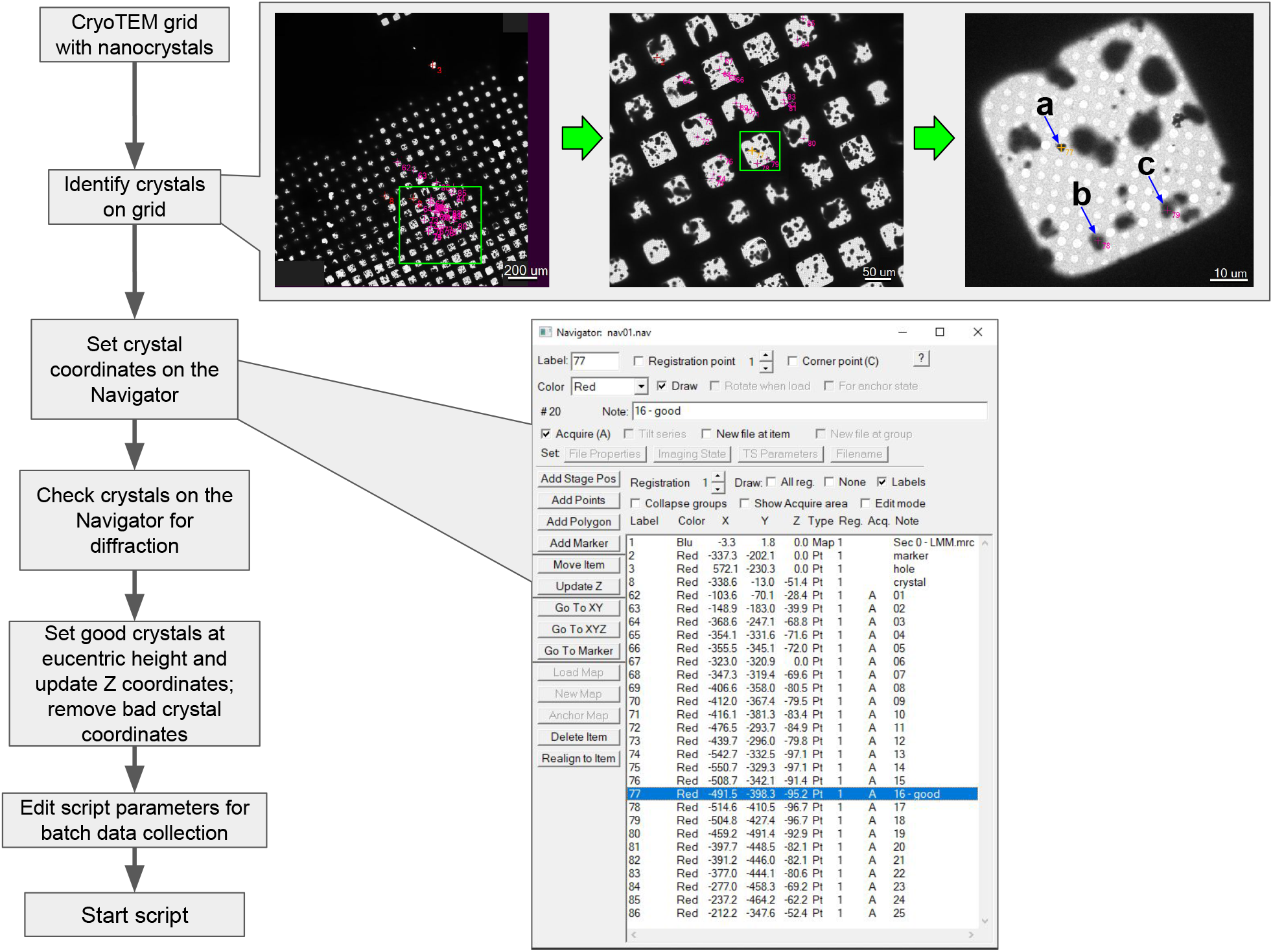
**SerialEM automation workflow for MicroED**. A cryoTEM grid containing protein nanocrystals is examined via a whole-grid atlas collected at low magnification. Crystal areas are then identified from this atlas and added into the SerialEM Navigator queue for subsequent diffraction screening. Diffracting crystals on this list are selected for subsequent MicroED data collection in batch, **a, b, c**, Crystals selected for diffraction screening.

**Figure 2.**
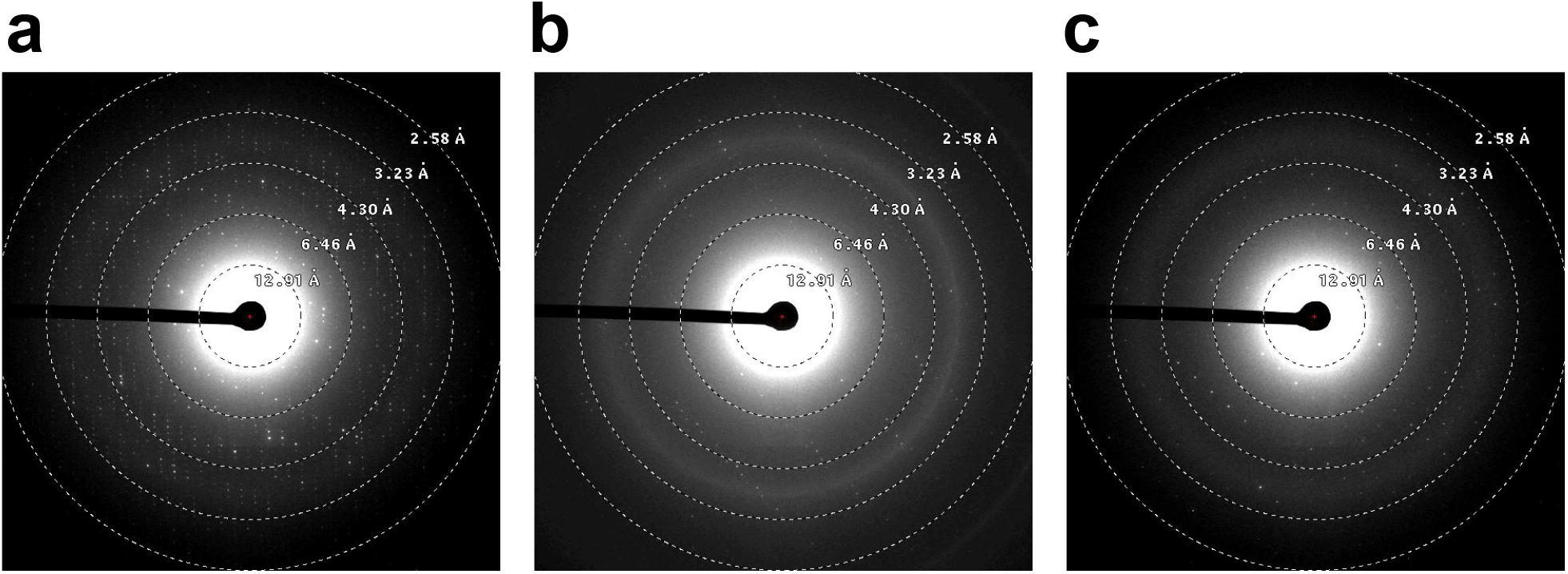
**Electron diffraction screening of selected crystals. a**,**b**,**c**, Diffraction patterns of crystals **a**,**b**,**c** in **Figure 1**.

Our pipeline coordinates all activities (microscope, crystal, and camera) to automate data collection. The operator inputs the desired rotation angle span (a range from −70° to +70° at maximum), rotation speed (degrees per second), exposure time per image frame (seconds), and destination directory for the data output; these are applied to all crystals on a job list. Data collection then proceeds in batch, where the SerialEM script directs the microscope sequentially through the list of selected crystals, one crystal at a time. SerialEM synchronizes crystal rotation and the start of data collection, such that the crystal is rotating at the desired, constant rate when the first frame is recorded. The entire rotation range is recorded as set, then camera recording stops, the stage rotation stops, and the data is saved. SerialEM then continues to the next crystal location in the job list. The relevant metadata, such as rotation rate and camera configuration, are automatically stored with the diffraction images in directories on the filesystem of the data collection host. These output files need to be converted to formats readable by typical crystallography integration packages such as DIALS^18^, iMosflm/MOSFLM^19^, or XDS^20^ using our conversion software^7^, which is freely available from https://cryoem.ucla.edu/pages/MicroED. The **Supplementary Protocol** contains a step-by-step procedure of SerialEM setup and instructions for MicroED data collection.

This work lays the foundation for automating MicroED data collection by using software to control the repetitive and intricate tasks involved, allowing the experimenter to spend more time on those aspects of structure determination where computer-controlled procedures are not yet feasible, such as crystal optimization and TEM grid preparation. Our procedures may be further developed to automatically detect crystals on a grid and determine optimal data collection parameters as is done at synchrotrons for X-ray crystallography (*e.g*., BEST^21^, STRATEGY^22^). Currently SerialEM only has access to the necessary microscope commands on Thermo Fisher systems; work on compatibility with JEOL microscopes is ongoing and support for other microscopes and cameras is anticipated soon.

Interest in cryoEM methods for biological investigation has recently surged as developments in hardware (increasingly stable electron optics, in-microscope robotic multiple-specimen holders allowing longer contamination-free storage times, and fast cameras allowing new ways of data collection) and software (automation for data collection and analysis) have opened cryoEM to the wider community. We show here that MicroED data can be collected in a consistent and automated way using the open-source software application SerialEM with minimal user intervention. This level of automation for MicroED simplifies the data collection experience for the end-user and makes MicroED accessible to laboratories and facilities without prior extensive experience. Depending on the type of camera used and the speed of rotation, data collection can take as little as 1 minute per crystal for a full 140° dataset that is often sufficient for structure determination. Under such a regime, data from more than 300 crystals can be collected autonomously overnight, bringing MicroED to the X-ray crystallography timescale and enabling analyses of big datasets that could be used for time-resolved studies, drug discovery, and protein dynamics. Thus, large-scale automation represents the next phase of MicroED studies.

## Methods

### Crystal growth

The fungal serine protease Proteinase K (*E. album*) from Sigma-Aldrich (St. Louis, MO, USA) was prepared by combining 3 mL of protein solution (50 mg mL^-1^) with 3 mL of precipitant solution (1.0–1.3 M ammonium sulfate, 0.1 M Tris @ pH 8.0)^15^ and set in a 24-well tray as 4-µL hanging drops.

### CryoEM sample preparation

3 µL of proteinase K solution was pipetted onto a Quantifoil R 2/2 Cu 300-mesh grid (Quantifoil Micro Tools GmbH, Großlöbichau, Germany) which had been pretreated with glow-discharge plasma at 15 mA for 30 s (PELCO easiGlow, Ted Pella Inc., Redding, CA, USA) and blotted onto 595 filter paper (Ted Pella Inc.) at a blot force setting of 5 for 10 s and plunge-frozen into liquid ethane using a Thermo Fisher Vitrobot Mark IV (Thermo Fisher Scientific, Hillsboro, OR) with a chamber environment of 22°C and 100% humidity.

### TEM configuration

Vitrified grids were examined with a Thermo Fisher Talos Arctica TEM operated at 200 kV and fitted with a Thermo Fisher CetaD scintillated CMOS 4k × 4k camera. The specimen was kept at ~100 K in the microscope. Low-magnification imaging for whole-grid crystal searching/montaging was done at a microscope magnification of 155×. TEM diffraction beam settings were saved into the SerialEM Low Dose Control for low-dose diffraction imaging^14^ and coherent-beam diffraction for data collection. In this Low Dose Control, the diffraction beam used for data collection was set to an electron dose of 0.01 e^-^ Å^-2^ s^-1^ and saved as the “Record mode” beam. The “View mode” beam setting was used for local (gridsquare) area imaging of crystals for screening on the FluCam (phosphor screen) of the Arctica TEM, and was set to an electron dose rate of ~1 x 10^-6^ e^-^ Å^-2^ s^-1^. A diffraction-defocus offset was applied to view the crystal using the SerialEM Low Dose Control’s defocus “offset for View mode” setting.

### Microcrystal screening

Grids were searched for thin ice and microcrystals first using the low magnification setting (155×). This initial search either took place manually, by observing the FluCam and moving the stage, or via the SerialEM Full Montage function, where a series of images are collected at the low magnification setting during a rasterization of the stage across the entire grid surface. Crystal stage coordinates were identified and saved to the SerialEM Navigator for later diffraction testing. To identify crystals suitable for data collection in an automated way, we set the Record mode camera parameters to collect a 4-second exposure. This uses the “Record Mode” diffraction beam setting as mentioned above for crystal exposure. The selected area (SA) aperture was inserted, the size of which was set to accommodate the dimensions of the largest crystal chosen for diffraction. We then used the “Acquire at Points” function to acquire diffraction pattern exposures at each previously-saved crystal coordinate. For those crystals which showed protein diffraction, the position was revisited to set the crystal at eucentric height using the microscope controls, then queued in the Navigator for batch MicroED data collection.

### MicroED data collection

Microcrystal coordinates prepared for data collection were set to “Acquire” in the SerialEM Navigator in the previous section. Before starting, the parameters for data collection, particularly output directory, rotation speed, angular range, frame exposure time, and frame binning, were configured in the CRmov script for this batch run. A detailed protocol for SerialEM-automated MicroED data collection and the CRmov script are available in the **Supplementary Protocol**.

## Code availability

The CRmov script and documentation are available in the **Supplementary Protocol**.

## Acknowledgments

We are grateful to David Mastronarde for feature support and technical assistance with SerialEM. This research was supported by the NIH/NCI Cancer Center Support Grant P30 CA008748 to Memorial Sloan Kettering Cancer Center. The Gonen lab is supported by funds from the Howard Hughes Medical Institute.

## Author contributions

M.J.d.l.C. and M.W.M. collected MicroED data. J.H. wrote data conversion software and processed data. M.W.M. processed data. M.J.d.l.C. conceived and designed the research, wrote the SerialEM data collection script, and prepared figures. T.G. coordinated the research. M.J.d.l.C. and T.G. wrote the manuscript with contributions from all authors.

## Competing Financial Interests

The authors declare no competing financial interests.

## A procedure for collecting MicroED data using SerialEM

M. Jason de la Cruz^1,2,*^

Michael W. Martynowycz^3^

Johan Hattne^3^

Tamir Gonen^1,3,*^

**Affiliations**

^1^ Janelia Research Campus, Howard Hughes Medical Institute, Ashburn, VA, USA

^2^ Structural Biology Program, Sloan Kettering Institute, Memorial Sloan Kettering Cancer Center, New York, NY, USA

^3^ Howard Hughes Medical Institute and Departments of Biological Chemistry and Physiology, David Geffen School of Medicine, University of California, Los Angeles, Los Angeles, CA, USA

^*^ Corresponding authors: M.J.d.l.C.; T.G.

### Abstract

We present a step-by-step procedure to automate microcrystal electron diffraction (MicroED) data collection of protein microcrystals using a script developed for use with the SerialEM software package on Thermo Fisher transmission electron microscopes (TEMs). Analogous to the way single-particle analysis (SPA) data is commonly collected via SerialEM, the script described here enables automated MicroED data collection from a sequential list of prepared crystal coordinates. Detailed steps for MicroED diffraction beam setup for Low Dose Mode in SerialEM are also outlined. Apart from the initial microscope and software setup, the timing for each step depends on several factors: 1) the quality of the crystal sample, 2) the number of crystals screened and selected for data collection, and 3) MicroED data collection parameters such as exposure time per frame and range of the angular sweep.

### Introduction

This protocol describes the setup for continuous-rotation MicroED^1–10^ crystal screening and automated multiple-crystal batch data collection using SerialEM^11^. The script “CRmov” described here will work with the Tecnai, Titan/Krios, and Talos series microscopes available from Thermo Fisher Scientific. Note that we use the phosphor screen for diffraction setup and screening, not the primary imaging detector; however, it may be desirable to use the detector for samples with very weak diffraction. If you do use the imaging device for these cases, do so with care and caution, as a focused high-energy direct beam may harm the detector. Also, we do not mention when to open or close the column valves, or whether to use a beamstop in this procedure; it is up to the user to use best practices when controlling dose to the sample and imaging detector. Our practice is to keep column valves closed or pre-specimen shutter inserted (beam blanker on) until absolutely necessary to avoid accumulating electron dose on the grid sample. We generally use a beamstop to prevent the direct beam from striking the detector surface.

This procedure assumes that the TEM has been optimally aligned for imaging in the SA lens magnification range, and SerialEM has been installed with proper microscope and camera communications as well as necessary stage and beam shift calibrations. It also assumes that the operator has a basic understanding of SerialEM, including the use of the Low Dose Control, the Navigator, and Scripts. We recommend the user to be familiar with the protocols for MicroED^3,6^ prior to setting up SerialEM.

### Equipment

#### Microscopy

- Thermo Fisher Scientific (formerly FEI Company, Philips Electron Optics) transmission electron microscope (TEM) of the Tecnai, Titan/Krios, or Talos series (with computer-controlled and driven specimen stage)
- Detector, usually a camera employing charge-coupled device (CCD) or complementary metal-oxide semiconductor (CMOS) imaging sensor, to capture and digitize the image formed from transmitted electrons produced by the TEM; either scintillated or non-scintillated/direct electron detection. The electron detector should be compatible with SerialEM; refer to the SerialEM website: http://bio3d.colorado.edu/SerialEM.
- Sufficient disk space for data. This depends on detector resolution, binning, and number of frames collected per dataset movie. For example, a full 4K × 4K unbinned dataset at 8 s per exposure with a rotation sweep of +30° to −30° at a rotation speed of ~0.18° s^-1^ is typically ~2 GB in size using a Thermo Fisher Talos Arctica with a Thermo Fisher CetaD camera.

#### Software

- SerialEM (version 3.7.0 beta10 64-bit or later, downloadable from http://bio3d.colorado.edu/ftp/SerialEM)
- SerialEM script: CRmov (see **Supplementary Document**)

#### Procedure

A. **Configure SerialEM Low Dose Mode for Diffraction**. In the “Low Dose Control” window (Figure 1), the Low Dose Mode is enabled by checking its box. Of the five possible areas that can be set, we are only interested in two for diffraction: View and Record. The SerialEM Record mode is configured with the beam set to diffraction. The View mode beam should be set to be defocused (diffraction defocus) from the Record mode beam, for low-dose “imaging” in diffraction mode (*i.e*., finding the crystal using a spread diffraction beam)^12^. Because View mode is an offset to the Record mode, we will set up the beam for diffraction data collection first for Record mode, then set the diffraction-focus offset for View mode. The Record mode beam is set up in a similar way to “exposure mode” as previously described^6^.
  1. **Record mode: Set the diffraction beam**. Enable Low Dose Mode, and make sure that “Continuous update” is checked. For “Go to area / show when screen down” choose “Rec.”. Set electron optics to diffraction mode by pressing the “Diffraction” button on the microscope console panel.
  2. On the microscope, make sure that you are in a beam-transparent area on the grid, preferably on a thin sample. Set camera length and condenser lens beam illumination for the dose that you require for the crystal (*i.e*., C1/spot size and C2 intensity or illuminated area).
  3. Insert beamstop and center the target area with the detector as necessary/possible. **Warning:** Please be advised that the next step involves focusing the beam to a coherent spot; you must take appropriate precautions to protect your imaging detector from excessive radiation before continuing.
  4. Please see warning in step **A3** above before proceeding. Use the focus knob in diffraction mode (diffraction focus) to condense the beam to a coherent spot (smallest diameter), then move it behind the beamstop using diffraction shift. This beam is now set up in the proper diffraction condition for data collection. To save the beam setting in SerialEM, uncheck the “Continuous update” box.
  5. **Tip 1:** Make sure that the objective aperture is in the “out” position.
  6. **Tip 2:** You may need to adjust the diffraction stigmator as necessary to ensure the beam is round; this should be done using the microscope controls.
  7. **Tip 3:** If the beam is sweeping as you move through crossover, you must center the condenser aperture(s).
  8. **Tip 4:** The ideal time for centering the SA (selected area) aperture is immediately following setup of the diffraction beam for screening/data collection.
  9. **Tip 5:** All illumination settings, when adjusting for a particular dose, should be done with sample retracted or otherwise not obstructing the beam (*i.e*., imaging in a hole/vacuum) for an accurate reading of the dose measurement.
  10. **View mode: Set beam for crystal search**. Once the Record mode has been set up, you can simply copy these settings to View mode as a base setting to which a defocus offset will be applied.
  11. Make sure you are still in Record Mode with Low Dose Mode enabled, and screen is down. Continuing from step **A4** above, under “Copy current area mag & beam to”, click the “V” button. This copies the current Record mode settings to View mode.
  12. Now go to View mode with the screen down: chose “Vie.” under “Go to area / show when screen down”. The mode changes but the beam parameters on the microscope should remain the same as Record mode.
  13. Enable “Continuous update” so you can change the View mode beam parameters.
  14. Change condenser lens beam illumination (spot size and C2 intensity/illuminated area) to desired values for low dose imaging. Because lens hysteresis can affect beam stability, it is advised to keep spot size the same for View mode and only vary C2 intensity/illuminated area to lower the dose for search (see also step **A16** below). On Thermo Fisher TEMs this can be achieved by increasing C2 intensity/illuminated area from the Record mode beam setting.
  15. In order to defocus the beam via diffraction focus, use the up/down arrow buttons under “Offsets for View” in the Low Dose Control. There is a limited range and fixed step size for defocus offset using this method, so beam edges and extreme pincushion distortion may be observed; however you should still be able to see your crystal. When satisfied with the defocused-diffraction image in View mode, save the beam setting by unchecking the “Continuous update” box.
  16. **Tip 1:** It is possible to reduce distortion in the View mode defocused-diffraction beam by changing defocus, camera length, spot size, and beam intensity as necessary. However, due to hysteresis in the magnetic lenses, it is best to minimize changes from the Record mode beam as much as possible for beam stability. We recommend altering only C2 intensity/illuminated area and diffraction defocus settings, then changing other parameters as necessary.
  17. **Tip 2:** All illumination settings, when adjusting for a particular dose, should be done with sample retracted or otherwise not obstructing the beam (*i.e*., imaging in a hole/vacuum) for an accurate reading of the dose measurement.
  18. **Center the SA aperture**. In order to select a single crystal for diffraction, the appropriate SA aperture size should be selected to limit diffraction from the area surrounding the crystal. It is appropriate to center the aperture when the data collection beam is being set up, since the aperture must be aligned to the beam’s optical axis.
  19. To make centering easier, place a sample with a landmark area (*e.g*., a crystal or hole pattern) in the path of the beam.
  20. With the screen down, make sure that the microscope is set to Low Dose Record mode as set above, with “Go to area / show when screen down” set to “Rec.”. Leave the “Continuous update” box unchecked, as we do not want to alter the setting for data collection.
  21. Ensure that the beam is centered on the beamstop/detector area as necessary, and that it is round and coherent.
  22. Defocus the beam using diffraction focus via the microscope control. If there is a sample in the beam, find a landmark on the grid, then center that landmark using the stage, so that the landmark appears in the middle of the beam.
  23. Insert the SA aperture and center it with the beam, or if a sample in the beam had been previously centered, center the aperture with the landmark.
  24. Once the aperture is centered, you can continue with screening or data collection. Switching to View and back to Record mode, with the screen down, should return the beam to the previous settings for each mode, as long as you did not check the “Continuous update” box during the SA aperture alignment. Otherwise, if the Record mode beam is not as expected, follow steps **A1-4** above to set the diffraction beam.
B. **Crystal search and screen using SerialEM Low Dose Mode, and queue for data collection**. Two approaches for local (gridsquare-level) search for crystals are described below using the View and Record beam modes of the SerialEM Low Dose Mode as set up above. Identified crystals ready for data collection are then added to a job list in SerialEM known as the Navigator.
  1. **Manual search**. A manual search for crystals may be done as previously described^12^ by using defocused-diffraction imaging to find and center crystals of interest, testing for diffraction (Record mode), then setting the crystal to eucentric height for data collection.
    a. **Set up SerialEM Low Dose View mode for live imaging**. Adjust the camera parameters for View mode to a short exposure setting (*e.g*., 0.5 s), and enable continuous acquisition mode. You can now use the “View” camera button in the “Camera & Script Controls” for live imaging in SerialEM. Remember to return the acquisition mode to “Single Image” when done with live imaging.
    b. **Search for crystals using the phosphor screen**. Make sure that the microscope is set to Low Dose View mode as set above, with “Go to area / show when screen down” set to “Vie.”. Then put the viewing screen down. If you cannot see the beam, make sure that it is unblanked.
    c. Use the microscope console stage controls to search for crystals using detector-assisted live imaging (step **B1a** above) or phosphor screen (step **B1b**).
    d. **Center a crystal in the SA aperture**. Make sure that the crystal is in view, then insert the SA aperture. Using the microscope stage controls, move the stage so that the crystal is centered in the beam, which is now limited by the SA aperture. You may need to insert and remove the SA aperture as necessary to locate the crystal.
    e. **Test a centered crystal for diffraction**. If using detector-assisted live imaging, use the “STOP” button under the “Camera & Script Controls” to stop live imaging. If using the phosphor screen to search for crystals, lift the screen and cover viewing window (if applicable).
    f. Adjust the camera parameters for Record mode to your desired exposure setting (*e.g*., 5 s) for diffraction mode.
    g. Insert beamstop (recommended).
    h. Press the “Record” button in the “Camera & Script Controls” section. This will switch the microscope to the previously-set diffraction settings in Record mode (steps **A1-4**), then expose the crystal at the exposure time set in step **B1f** above. The image may be saved to disk if desired using the options in the SerialEM File menu. If a diffracted crystal is satisfactory for data collection, *i.e*., spot intensities are sufficiently strong, continue to step **B1i** below. To test additional crystals without adding the current crystal position to the data collection queue, repeat the process from steps **B1c-h**.
    i. **Set crystal position to eucentric height**. Insert the SA aperture and ensure that the crystal is centered while in Low Dose View mode (either put the screen down in View mode, as described in step **B1b**, or use View mode imaging as in step **B1a**). If the crystal is not centered at this point, follow step **B1d** above to center it.
    j. While viewing the crystal, tilt the stage from one end of the range to the other and make sure that the crystal does not move away, or moves minimally from, the center of the aperture.
    k. Use the Z-height adjustment buttons on the microscope control panel to correct the eucentricity of the crystal within the tilt range.
    l. **Set a crystal coordinate on the SerialEM Navigator**. Once the crystal has been centered in three dimensions, it is ready for data collection. Open the Navigator if it is not already open.
    m. Add the current crystal position to the Navigator by pressing “Add Stage Pos” in the Navigator window (Figure 2).
    n. Set the Navigator point to “Acquire”.
    o. Repeat the steps **B1c-n** above for the next crystal.
    p. After your crystal positions are set, and SerialEM is properly set to run CRmov (steps **C1-3** below), you may edit the CRmov script to adjust batch collection parameters (step **C4** below), then set up and run the batch data collection as specified in step **C6** below.
  2. **Map-assisted search**. This allows offline searching for crystals using a low-magnification atlas of the entire grid. This procedure assumes that the beam image center (image shift) at low magnification is aligned to, or close to, the optical axis in the microscope. Otherwise, you will not be able to target the proper gridsquare/crystal in diffraction mode; or you must apply an offset to the crystal in View mode for proper targeting.
    a. **Collect a whole-grid atlas**. Open the Navigator if it is not already open.
    b. Using the “Set up full montage” feature in the Navigator menu, collect a whole-grid atlas (also called “full montage” or “Low Mag Montage/LMM”) of the grid by using the “Setup full montage” function in the Navigator menu. Consult the SerialEM documentation for more information on setup.
    c. **Add crystal targets using the Navigator**. Use the Navigator and whole-grid atlas to add points of crystal areas you wish to target/check for diffraction. Use the “Add Points” button on the Navigator, then left-click on the atlas to add crystal targets; click “Stop Adding” when done. This step can optionally be done “offline” because you are only interacting with the collected map in this step, not the microscope. Also, external review and selection of crystal targets can be done externally on another computer using the “DUMMY” version of SerialEM if the Navigator file and associated whole-grid atlas file are saved. Refer to SerialEM documentation for instructions on setting up the “DUMMY” version of SerialEM.
    d. At this point you have two options to check selected crystals for diffraction: manually (crystals are diffracted one-by-one; continue to steps **B2e-l** below) or semi-automatically (crystals centered in SA aperture manually, then diffraction patterns are collected in batch; continue to steps **B2m-v** below).
    e. **Check each Navigator point for diffraction: manual method**. Make sure that View and Record modes are set for diffraction as described above in steps **A1-4** above.
    f. Choose a saved crystal coordinate in the Navigator and press “Go To XY”. The microscope will move the stage to the selected target.
    g. **Center a crystal in the SA aperture**. Use detector-assisted live imaging (step **B1a** above) or the phosphor screen (step **B1b**) to make sure that the crystal is in view, then insert the SA aperture. Using the microscope stage controls, move the stage so that the crystal is centered in the beam, which is now limited by the SA aperture. You may need to insert and remove the SA aperture as necessary to locate the crystal.
    h. **Test a centered crystal for diffraction**. If using detector-assisted live imaging, use the “STOP” button under the “Camera & Script Controls” to stop live imaging. If using the phosphor screen to search for crystals, lift the screen and cover viewing window (if applicable).
    i. Adjust the camera parameters for Record mode to your desired exposure setting (*e.g*., 5 s) for diffraction mode.
    j. Insert beamstop (recommended).
    k. Press the “Record” button in the “Camera & Script Controls” section. This will switch the microscope to the previously-set diffraction settings in Record mode (steps **A1-4**), then expose the crystal at the exposure time set in step **B2i** above. The image may be saved to disk if desired using the options in the SerialEM File menu. If a diffracted crystal is satisfactory for data collection, *i.e*., spot intensities are sufficiently strong, then first make sure that the current crystal coordinate (from step **B2f**) is highlighted. Delete this coordinate from the Navigator by pressing “Delete Item”. Then, set the crystal position to eucentric height as described in steps **B1i-k**. Next, press “Add Stage Pos” to add this crystal’s updated (*x,y,z*) coordinates to a new Navigator point. Set this point to “Acquire” in the Navigator. Repeat steps **B2e-k** for each crystal in the Navigator. When done, continue to step **B2l** below.
    l. After your crystal positions are set, and SerialEM is properly set to run CRmov (steps **C1-3** below), you may edit the CRmov script to adjust batch collection parameters (step **C4** below), then set up and run the batch data collection as specified in step **C6**.
    m. **Check each Navigator point for diffraction: batch mode**. Continuing from step **B2d** above, make sure that View and Record modes are set for diffraction as described above in steps **A1-4**.
    n. Choose a saved crystal coordinate in the Navigator and press “Go To XY”. The microscope will move the stage to the selected target.
    o. **Center a crystal in the SA aperture**. Use detector-assisted live imaging (step **B1a** above) or the phosphor screen (step **B1b**) to make sure that the crystal is in view, then insert the SA aperture. Using the microscope stage controls, move the stage so that the crystal is centered in the beam, which is now limited by the SA aperture. You may need to insert and remove the SA aperture as necessary to locate the crystal.
    p. Make sure that the current crystal coordinate (from step **B2n**) is highlighted. Delete this coordinate from the Navigator by pressing “Delete Item”. Then, press “Add Stage Pos” to add this crystal’s updated (*x,y,z*) coordinates to a new Navigator point. Repeat steps **B2m-p** for each crystal in the Navigator. When done, continue to step **B2q** below.
    q. In the Navigator, set the updated points you wish to test for diffraction (corresponding to the selected crystals) to “Acquire”.
    r. Adjust the camera parameters for Record mode to your desired exposure setting (*e.g*., 5 s) for diffraction mode.
    s. Open a new file to save images by going to “File > Open New…”.
    t. In the SerialEM menu, go to “Navigator > Acquire at Points…”. In the “Acquire at Items” window that appears (Figure 3), under “Primary Task”, choose “Just acquire and save image or montage”, then click “GO”. This will instruct SerialEM to move the microscope stage to each crystal position and collect a Record mode exposure using the exposure settings defined in step **B2r** above. The exposures are saved in the file in the order of the queue. After this task has finished, and as long as the file is open and active in SerialEM, you may review the exposures (diffraction patterns) in the file by choosing “File > Read”. For each crystal that displays satisfactory diffraction for MicroED data collection, *i.e*., spot intensities are sufficiently strong, set the crystal position to eucentric height as described in steps **B1i-k**, press “Update Z” in the Navigator, then set the corresponding Navigator point to “Acquire” (Figure 2).
    u. When done, you may close the file opened in step **B2s** by going to “File > Close”.
    v. After your crystal positions are set, and SerialEM is properly set to run CRmov (steps **C1-3** below), you may edit the CRmov script to adjust batch collection parameters (step **C4** below), then set up and run the batch data collection as specified in step **C6** below.
C. **Continuous-rotation MicroED data collection using CRmov**. A SerialEM script called “Continuous Rotation movie acquisition”, or CRmov, was developed to automatically tilt to the starting angle, start continuous stage rotation, and continuously collect exposures until the end angle, saving these to files within a specified directory on disk. The resulting movie can then be converted to a standard crystallography format using tools^3^ available at https://cryoem.ucla.edu/pages/MicroED for subsequent processing.
  1. **Install CRmov to SerialEM**. A file called CRmov.txt is included with this protocol. This file may be imported into the SerialEM Script Editing Window by opening an “Edit” slot under the Script menu. Choose an empty Script Editing Window (*e.g*., Script > Edit 1; see Figure 4). Click the “Load” button at the bottom of the window and choose the CRmov.txt file to open it. Alternatively, you can open CRmov.txt in a text editor such as Notepad and cut-and-paste the text into the Script Editing Window.
  2. **Troubleshooting note:** Please be careful when editing the scripts, as typographical mistakes may cause unintended operation. It is important to make sure that all instances of the equals sign (=) are padded with a single space on both sides.
  3. **SerialEM properties file prerequisite**. This script uses the BackgroundTilt command which requires a fourth network socket connection to the microscope computer for continuous rotation to run concurrent with other SerialEM tasks. Add the following entry to the properties file after defining the SocketServerIPif64 and SocketServerPortIf64 entries: “BackgroundSocketToFEI 1”.
  4. **Using CRmov: Set up**. This script is designed to be used in conjunction with the SerialEM Navigator, where a user would queue up crystal coordinates for the same data collection parameters for each crystal. Crystal coordinates which require different data collection parameters should be queued up separately for a later run. This section will describe settings that should be checked and changed as necessary for each batch data collection session. These settings are in the “CHANGEABLE SETTINGS” section at the top of the script as seen in Figure 4. It is important to note that all points set to Acquire during a single run will use the settings entered in the script. If you wish to use different values for certain data collection parameters, such as path to root directory, angular range, rotation speed, rotation direction, etc., for different crystal coordinates, you must run those points in a separate batch run. The CRmov.txt file included with this protocol is pre-populated with values for reference.
    a. **fullPath**. Full Windows path of the root directory where you want the MicroED datasets saved, with no trailing backslash; *e.g*., E:\Jason\20180719\20180720_ProtK_1B. For each crystal (*i.e*., each point in the Navigator set to “Acquire at points“), the IDOC metadata file (text file with extension .idoc) is created along with TIF images of the continuous-rotation dataset and will be saved in a separate subdirectory of the format “MmmDD_hh.mm.ss”, where Mmm is the abbreviation for month, DD for day, hh for hour, mm for minute, and ss for second; *e.g*., “Jul20_10.00.30”. Therefore, each crystal dataset is organized in subdirectories by time/date stamp.
    b. **rotationSpeed**. The constant speed at which the goniometer rotates during data collection. This number is a multiplier to the Compustage property in the TEMspy/TADui (Thermo Fisher) program that controls this setting. In our experience, the value 0.0032 typically corresponds to ~0.09° s^-1^.
    c. **angleStart**. Relative angle, from the goniometer’s zero position in the TEM, at which the rotation sweep starts.
    d. **angleEnd**. Relative angle, from the goniometer’s zero position in the TEM, at which the rotation sweep ends.
    e. **Direction**. This signals the script to use the proper routine for stopping the stage rotation. It is important that this parameter is set properly! For rotation in the positive direction, enter 1. For rotation in the negative direction, enter 0.
    f. **plusMinusRange**. This is the buffer range for the end angle for use by the algorithm used to stop the stage rotation. Because of goniometer and sensor imprecision, the stage may or may not stop exactly at the specified angleEnd. This parameter takes this into account by stopping stage rotation in this angular range abutting the angleEnd. The default value is 0.2, which should be good for most angular range/rotation speed regimes.
    g. **frameExposureTime**. Exposure time per image in the rotation dataset.
    h. **frameBinning**. Binning level of the saved exposures; a value of 1 indicates unbinned data and is the default value. i. **imagingMag**. Magnification, in TEM imaging mode, for scopes that require stage calibration in imaging mode. This is generally set to the mag setting when exiting out of diffraction mode. Default is 1250.
  5. **Using CRmov: Advanced settings**. The following parameters are preset settings that should work for most microscope/camera setups, and should not normally need to be changed.
    a. **currentCamera**. Camera to use, where the order is defined from the SerialEM properties file. Cameras are numbered from 1; if only one camera exists, this value should be 1.
    b. **recordMode**. Typically R for Record mode. This normally does not need to be changed, however, if there are problems setting/saving the diffraction spot it is possible to attempt using a different Low Dose Mode for data collection, *i.e*., F for Focus mode.
    c. **numberFrames**. This is the maximum number of frames for SerialEM’s continuous mode to collect before it stops. Ideally this parameter should be set to just past the tilt range; if set to before the end of the tilt range, data saving will stop before rotation completion. To be safe, the default value is set to a generous 400 to prevent premature recording end.
    d. **delayTime**. Delay for stage settling after tilting to angleStart. Default is set to half of the frameExposureTime, or $frameExposureTime / 2.
  6. **Using CRmov: Start a batch data collection run**. This process assumes that SerialEM is properly configured to run CRmov (steps **C1-3** above) and CRmov script parameters are set as necessary for the crystal coordinates in this batch run (steps **C4,5** above). The script will override settings in SerialEM Camera Parameters to collect data at the binning and exposure time set in the script as described above in step **C4**. The steps below follow from either step **B1p, B2l**, or **B2v** above, where you should already have crystal coordinates, at eucentric height and ready for continuous-rotation MicroED data collection, set to “Acquire” in the Navigator.
    a. Make sure that the beamstop is inserted (if necessary) before starting.
    b. Ensure that the **fullPath** parameter is set properly in the script. This entry should be a directory on the local computer (*i.e*., local to SerialEM); refer to step **C4a** above for more information.
    c. It is recommended to open the Log window (from the SerialEM taskbar menu: File > Open Log) to monitor progress and save as a reference for processing.
    d. In the SerialEM taskbar menu at the top of the screen, choose “Navigator > Acquire at Points…”. A dialog box titled “Acquire at Items” opens.
    e. In the “Acquire at Items” dialog window, in the “Primary Task” section, select “Run script” then in the drop-down menu beside it, choose the CRmov script. Refer to Figure 3 for reference. Click “GO” to start.
    f. For each crystal position (*i.e*., each point in the Navigator set to Acquire (A), the batch run will create a subdirectory within the specified **fullPath** (refer to step **C4a** above). The process creates sequential TIF files of continuous-rotation MicroED data and a text file with the filename corresponding to the Navigator Point/Label number and file extension .idoc. This file is the IDOC file containing metadata information for each frame collected.
    g. After data collection is complete, save the data in the Log window (File > Save Log…), and optionally save the Navigator file if needed.

**Figure 1.**
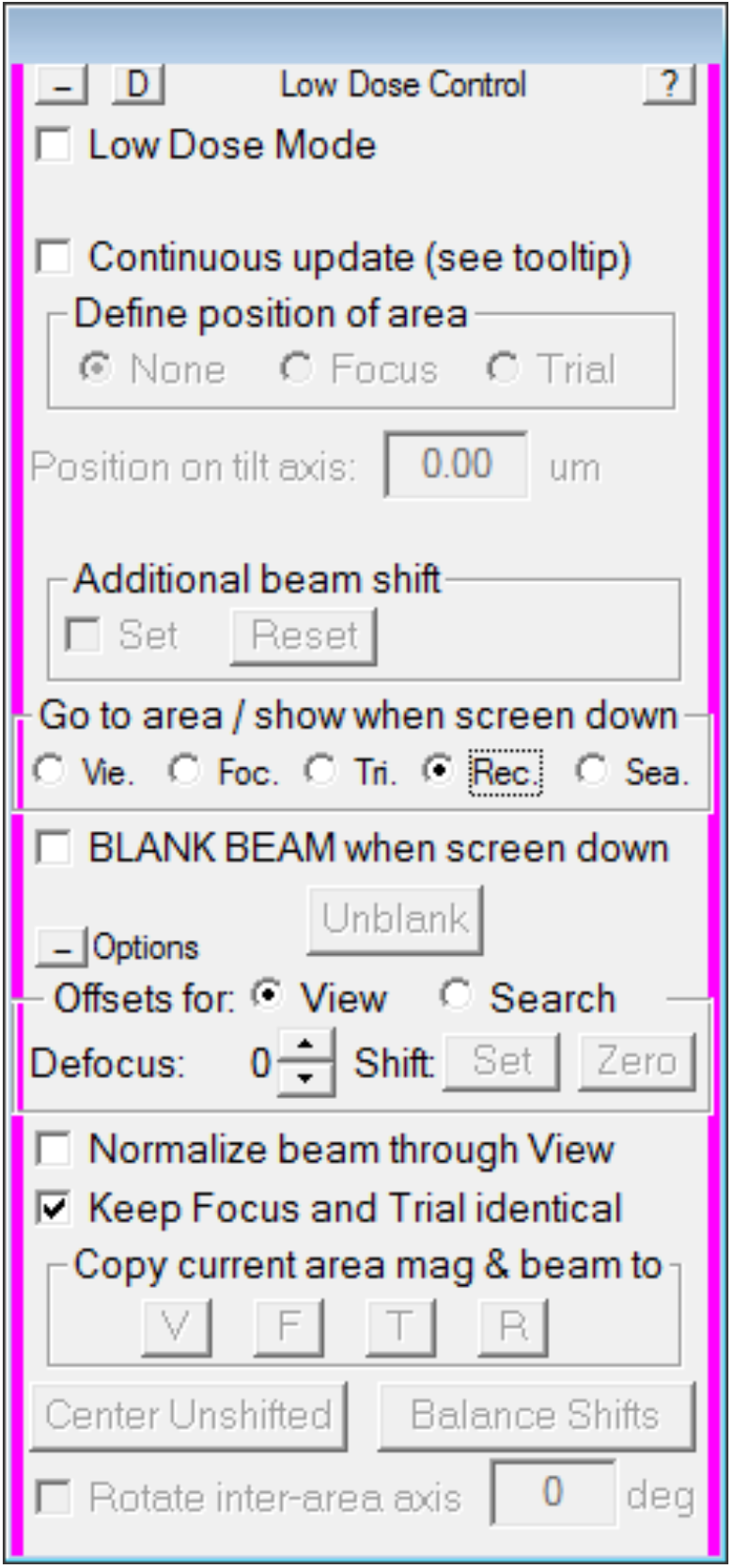
SerialEM Low Dose Control window.

**Figure 2.**
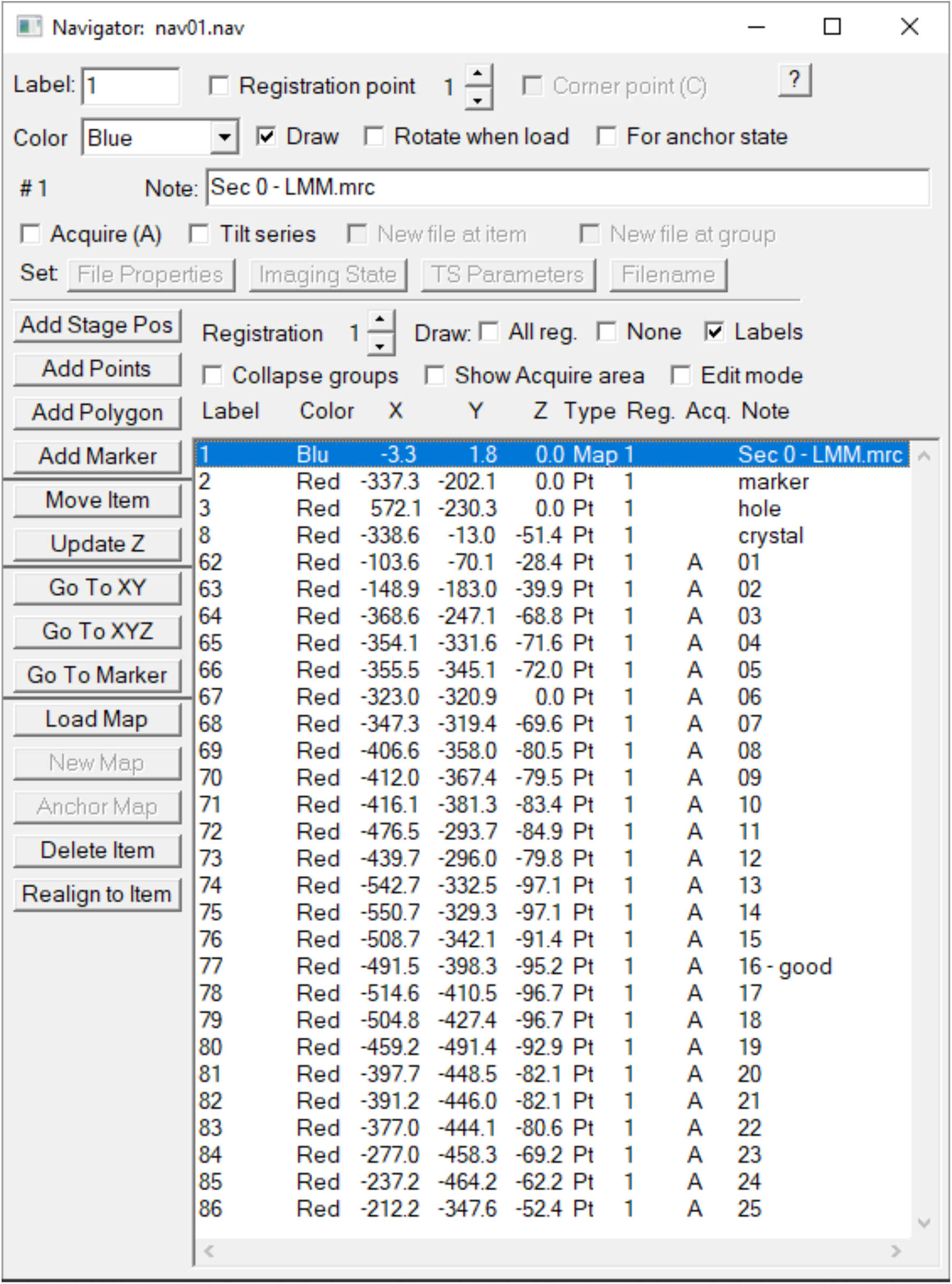
Navigator window with crystal points set to “Acquire”.

**Figure 3.**
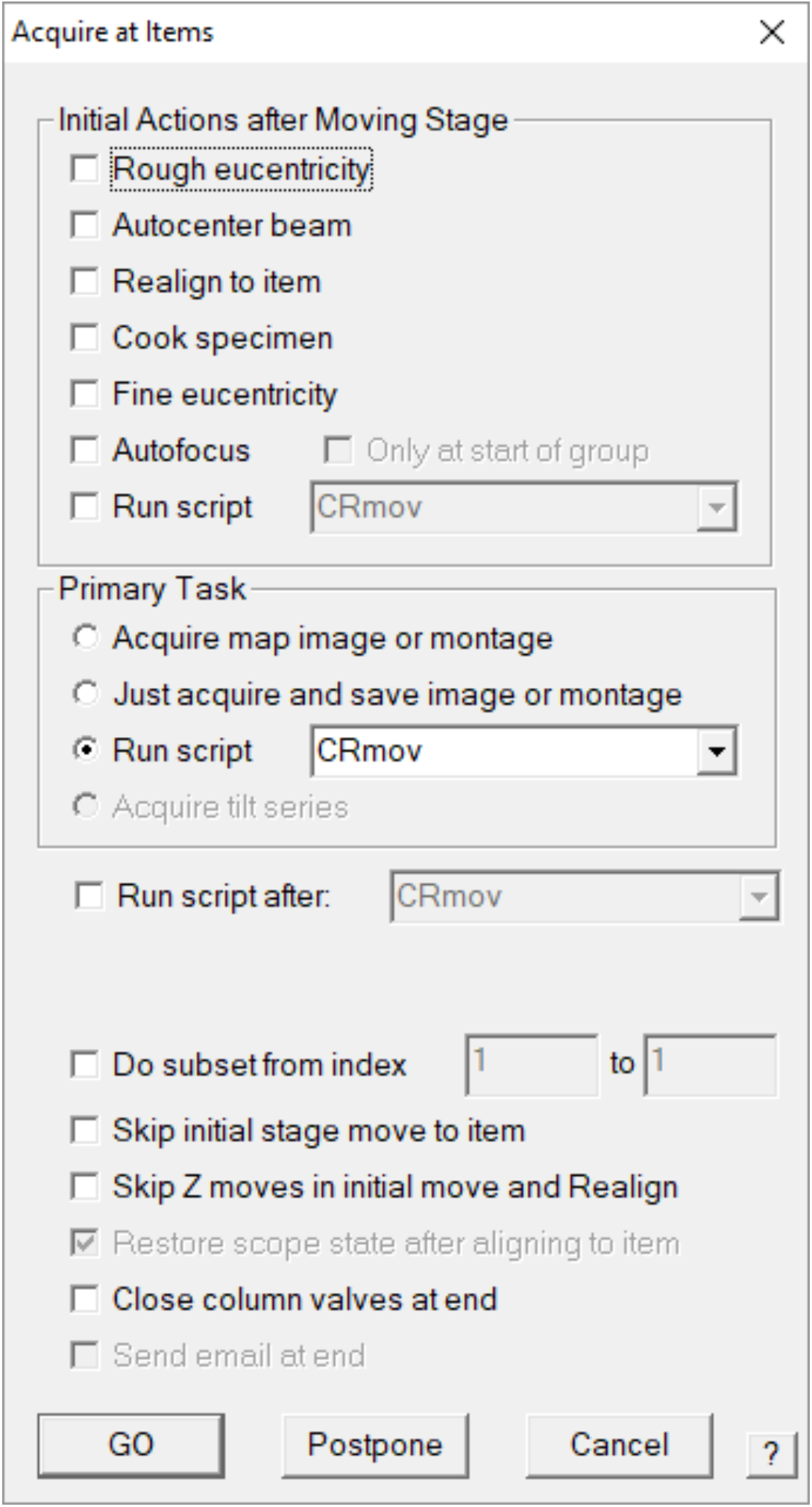
“Acquire at Items” dialog window.

**Figure 4.**
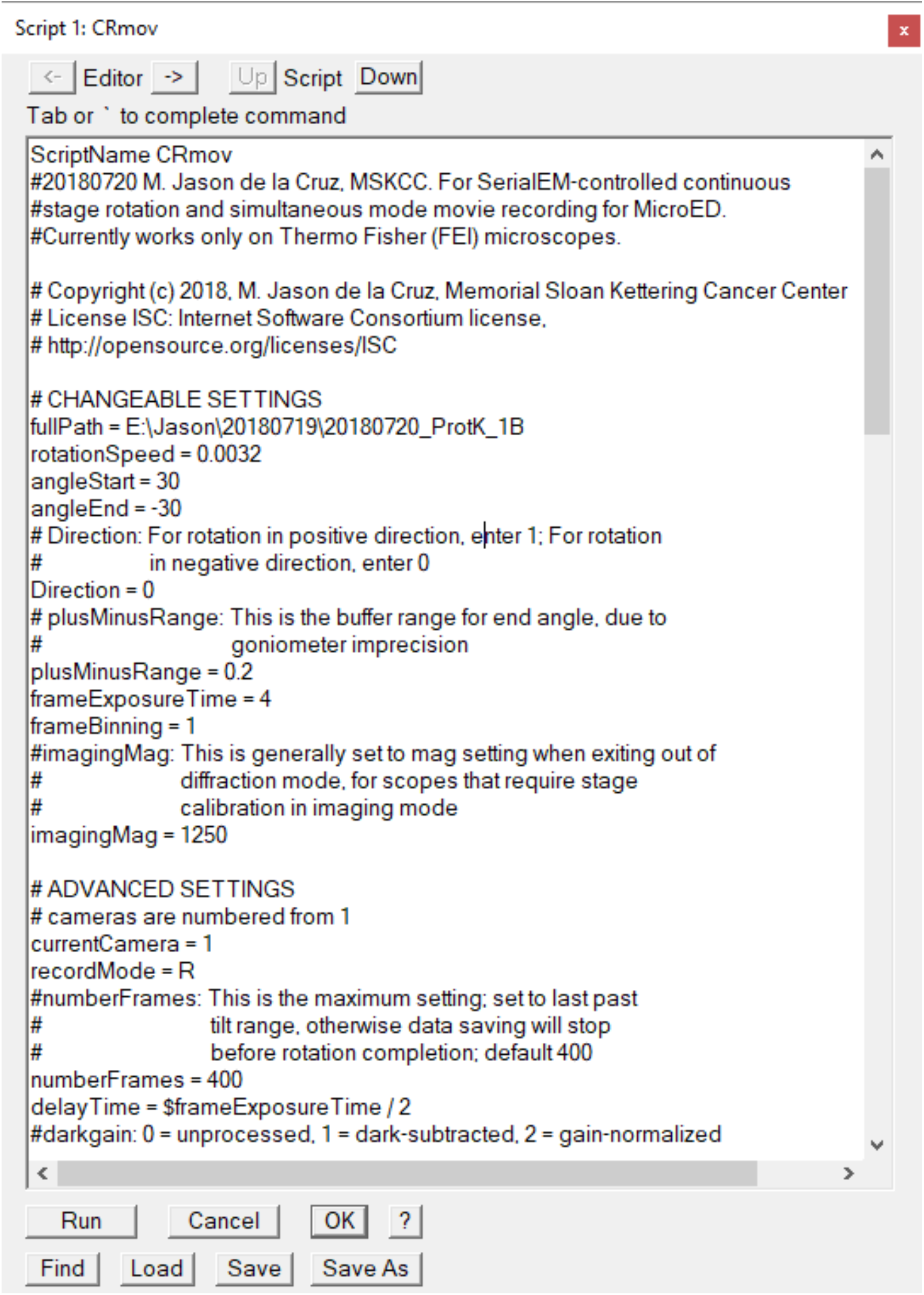
Script Editing Window showing the CRmov script loaded in the “Edit/Script 1” slot.

### Timing

Experiment timing depends on crystal screening and data collection parameters, therefore it may vary greatly. In our tests using a Thermo Fisher CetaD camera coupled to a Thermo Fisher Talos Arctica TEM, a single crystal dataset, collected using a rotation sweep of +30° to −30° at a rotation speed of 0.5° s^-1^, consisted of 200 frames of 4K × 4K data at 2 s per exposure and took 4 minutes to complete.

## Code availability

The CRmov script is available as a **Supplementary Document**.

## Acknowledgments

We thank David Mastronarde for feature support and technical assistance with SerialEM. This research was supported by the NIH/NCI Cancer Center Support Grant P30 CA008748 to Memorial Sloan Kettering Cancer Center. The Gonen laboratory is supported by funds from the Howard Hughes Medical Institute.

## Competing Interests

The authors declare no competing financial interests.

## Supplementary Document

CRmov.txt is a script for SerialEM which can be imported into SerialEM’s Script Editing Window. Updated versions of this script will be deposited to the SerialEM Script Repository located here: https://serialemscripts.nexperion.net.

### CRmov.txt

`ScriptName CRmov`

`# 20180720 M. Jason de la Cruz, MSKCC. For SerialEM-controlled continuous`

`# stage rotation and simultaneous mode movie recording for MicroED.`

`# Currently works only on Thermo Fisher (FEI) microscopes.`

`# Copyright (c) 2018, M. Jason de la Cruz, Memorial Sloan Kettering Cancer Center`

`# License ISC: Internet Software Consortium license,`

\# http://opensource.org/licenses/ISC

`# CHANGEABLE SETTINGS`

`fullPath = E:\Jason\20180719\20180720_ProtK_1B rotationSpeed = 0.0032 angleStart = 30 angleEnd = −30`

`# Direction: For rotation in positive direction, enter 1; For rotation`

`# in negative direction, enter 0 Direction = 0`

`# plusMinusRange: This is the buffer range for end angle, due to`

`# goniometer imprecision plusMinusRange = 0.2`

`frameExposureTime = 4`

`frameBinning = 1`

`#imagingMag: This is generally set to mag setting when exiting out of`

`# diffraction mode, for scopes that require stage`

`# calibration in imaging mode imagingMag = 1250`

\# ADVANCED SETTINGS

`# cameras are numbered from 1 currentCamera = 1 recordMode = R`

`# numberFrames: This is the maximum setting; set to last past`

`# tilt range, otherwise data saving will stop`

`# before rotation completion; default 400 numberFrames = 400`

`delayTime = $frameExposureTime / 2`

`# darkgain: 0 = unprocessed, 1 = dark-subtracted, 2 = gain-normalized darkgain = 2`

`echo ========================================`

`echo ===> Running CRmov …`

`echo ========================================`

`ProgramTimeStamps`

`SetDirectory $fullPath MakeDateTimeDir ReportNavItem SetNewFileType 1 ReportDirectory echo $navLabel`

`OpenNewFile $navLabel.idoc`

`ScreenUp SetLowDoseMode 1 GoToLowDoseArea $recordMode SetColumnOrGunValve 1`

`SelectCamera $currentCamera`

`CameraProperties`

`AddToAutodoc PhysicalPixel $reportedValue4`

`ReportCameraLength`

`AddToAutodoc CameraLength $reportedValue1`

`AddToAutodoc RotationRate $rotationSpeed`

`WriteAutodoc`

`SetExposure $recordMode $frameExposureTime 0`

`SetBinning $recordMode $frameBinning`

`SetCameraArea $recordMode F`

`SetProcessing $recordMode $darkgain`

`echo ===> Tilting to $angleStart degrees`

`TiltTo $angleStart`

`echo ===> Preparing beam and settling…`

`StopContinuous`

`GoToLowDoseArea $recordMode`

`echo Waiting for $delayTime seconds`

`Delay $delayTime sec`

`UseContinuousFrames 1`

`echo angleEnd is $angleEnd`

`echo rotationSpeed is $rotationSpeed`

`BackgroundTilt $angleEnd $rotationSpeed`

`Loop $numberFrames index`

`echo === START PROCESS FOR EXPOSURE $index of $numberFrames max frames ===`

`echo --- Resetting clock resetclock`

`echo Recording $frameExposureTime -second exposure`

`$recordMode`

`WaitForNextFrame`

`echo --- Saving exposure $index with initial angle`

`S`

`reportclock`

`ReportTiltAngle`

`CurrentAngle = $ReportedValue1`

`echo Current Tilt Angle is: $CurrentAngle`

`if $Direction == 0`

`negLimit = $angleEnd + $plusMinusRange`

`echo Specified tilt direction is negative with end angle stop at $negLimit degrees`

`if $CurrentAngle < $negLimit`

`break`

`else`

`continue`

`endif else`

`posLimit = $angleEnd - $plusMinusRange echo Specified tilt direction is positive with end angle stop at $posLimit degrees`

`if $CurrentAngle > $posLimit`

`break`

`else`

`continue`

`endif endif`

`EndLoop`

`echo ===> Resetting stage alpha to zero Delay 1 TiltTo 0`

`echo Acquisition complete`

`CloseFile`

`SetDirectory $fullPath`

`ReportDirectory`

`UseContinuousFrames 0`

`RestoreCameraSet $recordMode`

`GoToLowDoseArea R`

`SetColumnOrGunValve 0`

`SetLowDoseMode 0`

`Delay 1`

`SetMag $imagingMag`

`ProgramTimeStamps`

`echo ===> Scope standby`

